# Biowheel: interactive visualization and exploration of biomedical data

**DOI:** 10.1101/099739

**Authors:** Chenyue W. Hu, Alex J. Bisberg, Amina A. Qutub

## Abstract

We introduce Biowheel (https://biowheel.dibsvis.com/), a web-based award-winning data visualization tool, for exploring high-dimensional and heterogeneous biomedical data. Through interactive sorting and filtering of data, Biowheel enables researchers to quickly detect data outliers, evaluate data consistency, and discover mixed trends. Its interactive data presentation, visually-engaging design, and friendly user interface opens the door to easier, faster and better high-dimensional data interpretation for biomedical professionals with and without programming training.

---

How to effectively visualize and explore high-dimensional data remains an active field of research in biomedicine. Recent years have witnessed a fast expansion of measuring dimensions (e.g., number of genes, samples, time points) brought by advances in high-throughput omics and sensor technologies^1–3^. Meanwhile, the degree of heterogeneity observed in biomedical data of the same type is rapidly increasing, thanks to improved resolution in measurements^4^, the awareness of tumor heterogeneity^5^, and a growing interest in personalized medicine^6^. This unprecedented scale, diversity, and heterogeneity of biomedical data calls for developments of unconventional visualization methods and novel software tools to drive data interpretation^7^. To that end, the HPN-DREAM breast cancer network inference crowd-sourced data challenge^8^ in 2013 dedicated a sub-challenge to crowd-source visualization strategies for high-dimensional molecular time-course data sets in breast cancer.

Here, we present Biowheel, a data visualization tool created from the winning design of the HPN-DREAM visualization sub-challenge. The idea of Biowheel was inspired in part by the aesthetics of circos^9^, the utility of heatmaps^10^, and the powerful interactivity of web-based visualization frameworks^11^. Circular heatmaps, enabling end-to-end comparison, serve as the core design in Biowheel to visualize both numeric and categorical data. Differentiating its design from other applications, Biowheel is fully interactive and drives data interpretation through interactive display, filtering and sorting of the raw data. In addition, Biowheel frees biomedical researchers from programming, and speeds up the scientific discovery process with its easy-to-learn graphical user interface.

An example of visualizing high-dimensional molecular time-course data with Biowheel is shown in Figure 1A, using the main experimental breast cancer proteomics data set from the HPN-DREAM challenge. The original data set contains reverse phase protein array (RPPA) expression measurements of 45 phospho-proteins treated with 4 types of inhibitor and 8 types of stimulus at 7 post-stimulus time points in 4 cell lines. In this example, the time-course protein expression with 2 stimuli (EGF and FGF1) and 2 inhibitors (GSK690693 and PD173074) in the BT20 cell line is shown, where the time course starts from the innermost ring and projects outwards, and the proteins are represented as spokes. The specific types of inhibitor and stimulus being used in the experiment are denoted in the two outermost rings. A step-by-step guide of how to build this visualization in Biowheel is available in Supplementary Video 1.

**Figure 1.**
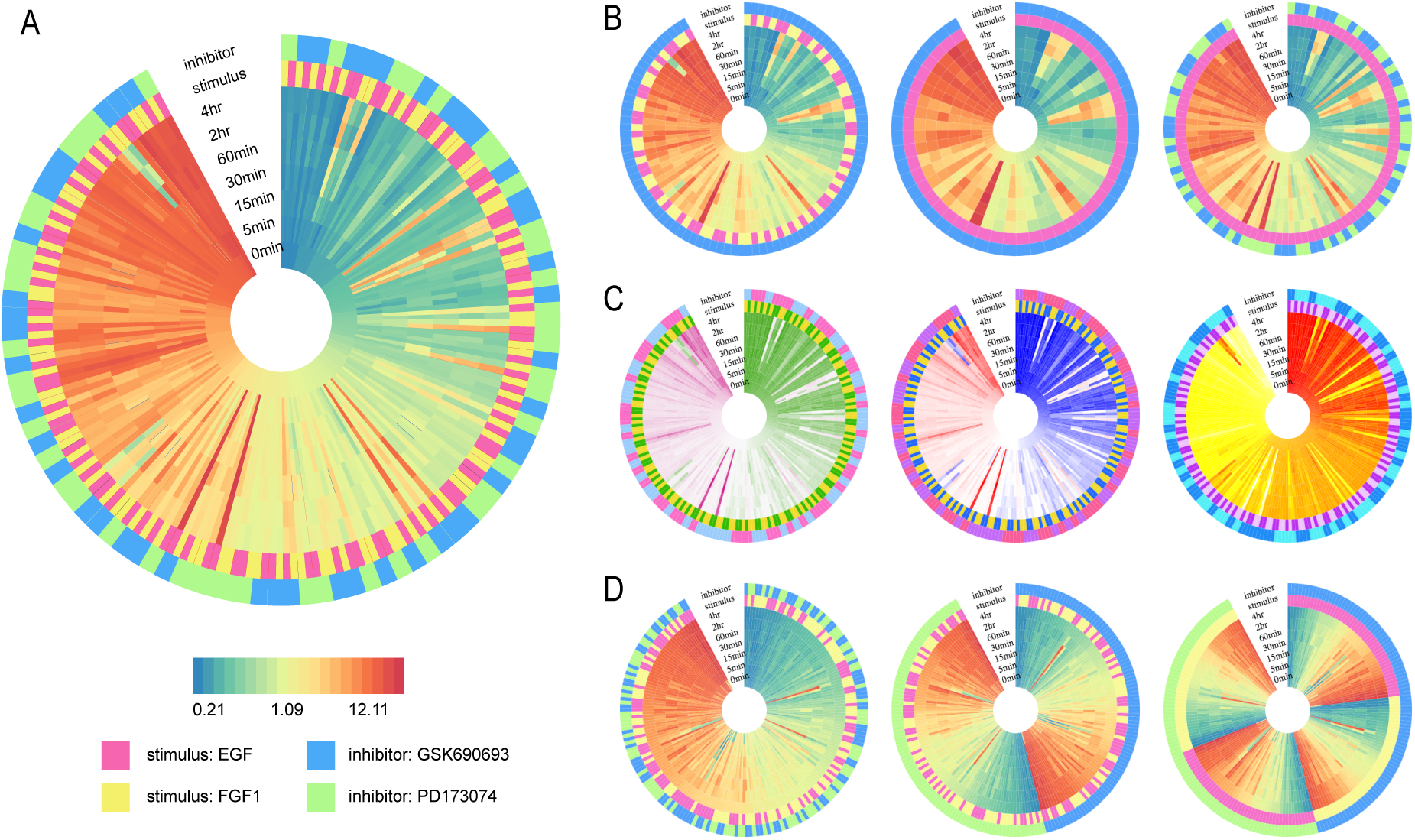
Assorted visualization of the HPN-DREAM challenge data in Biowheel: (A) time-course protein abundance under the inhibition of GSK690693 or PD173074 and the stimulation of EGF or FGF1 is shown, in which proteins are sorted by their expression levels at 0 min. (B) Three filtered subsets of (A) are generated by de-selecting samples inhibited by GSK690693 (left), by GSK690693 and FGF1 (middle), and by FGF1 (right). (C) Three coloring variations of (A) are created to demonstrate Biowheel’s aesthetic versatility. (D) Three spoke orderings of (A) are shown, in which spokes are sorted by expression at 4 hrs (left), by expression at 4 hrs and inhibitor (middle), and by expression at 4 hrs, inhibitor and stimulus (right). Supplementary Video 2 introduces all these features in a live demo.

Though initially proposed for visualizing time-course expression data, Biowheel can easily visualize any tabular data, independent of the data source and context. Each column from the tabular data will be visualized as a ring, and users are able to choose whether to represent it as a continuous variable, a categorical variable, or both. In the previous example of visualizing the HPN-DREAM challenge data (Figure 1A), expression levels at each time point are selected as a continuous variable, whereas experimental conditions are selected as categorical variables. Each row from the input tabular data is visualized as a spoke in Biowheel, and spokes that share the same characteristics can be excluded or included based on the user’s preference. An example of spoke selection is demonstrated in Figure 1B, where samples were excluded from the graph based on the inhibitor type, the stimulus type, or both. In contrast to the data filtering that we typically see in programs like Excel, Biowheel can filter data with multiple variables, and the selection is reflected instantly in the visualization.

In the spirit of heatmaps^12^, Biowheel presents data values in colors to empower visual interpretation. Continuous variables are mapped to a spectrum of colors (e.g. rainbow, heat, red-white-blue), and each level of categorical variables is represented by a unique color. Colors in every graphical element can be readily modified with clicks on the graph, thus achieving aesthetic versatility for the graph (Figure 1C). Besides intuitive color mappings, the numerical or categorical value of each data point can be simultaneously displayed on top of the graphical element to add more depth to the graph. The data display is therefore also interactive, with values popping up in a semi-transparent tooltip box when users mouse over the graph to inspect data points of interests.

Beyond its aesthetic visualization capabilities, Biowheel can facilitate data vetting and pattern discovery. The key feature enabling this is interactive sorting. Upon clicking any ring’s name in the graph, Biowheel will automatically sort (or unsort) all samples based on values of the corresponding variable and will reorder spokes instantaneously. Figure 1D illustrates three varied representations of the same data after sorting it by combinations of different variables. The sorting becomes nested once there are two or more variables being selected, and the hierarchy of the nested sorting is determined by the sequence of the selections.

The feature of nested sorting enables visual comparison between multiple data sources. For example, Biowheel can be used to visualize and compare dynamic responses of proteins in multiple experimental conditions. Using a subset of the HPN-DREAM challenge data (inhibitor: GSK690693; stimulus: EGF, FGF1), Figure 2A illustrates time-course protein expression data which is sorted by the stimulus type and protein expression at 0 min. Since the initial expression is the same for each protein across all stimulus conditions, the 45 proteins are arranged by the same order within the two sub-wheels divided by the stimulus. In this arrangement, the contrasting dynamic responses of proteins under EGF versus FGF1 stimulation appear immediately: a substantial portion of proteins that were of low or medium expression at 0 min were up-regulated when stimulated by EGF, but not when stimulated by FGF1. In addition, we can see a couple of cases in which a highly-expressed protein dropped its expression significantly after 60 minutes of FGF1 stimulation, which is not observed in the EGF stimulation experiments. These cases can also be interpreted as data outliers, which Biowheel is fully equipped to detect.

**Figure 2.**
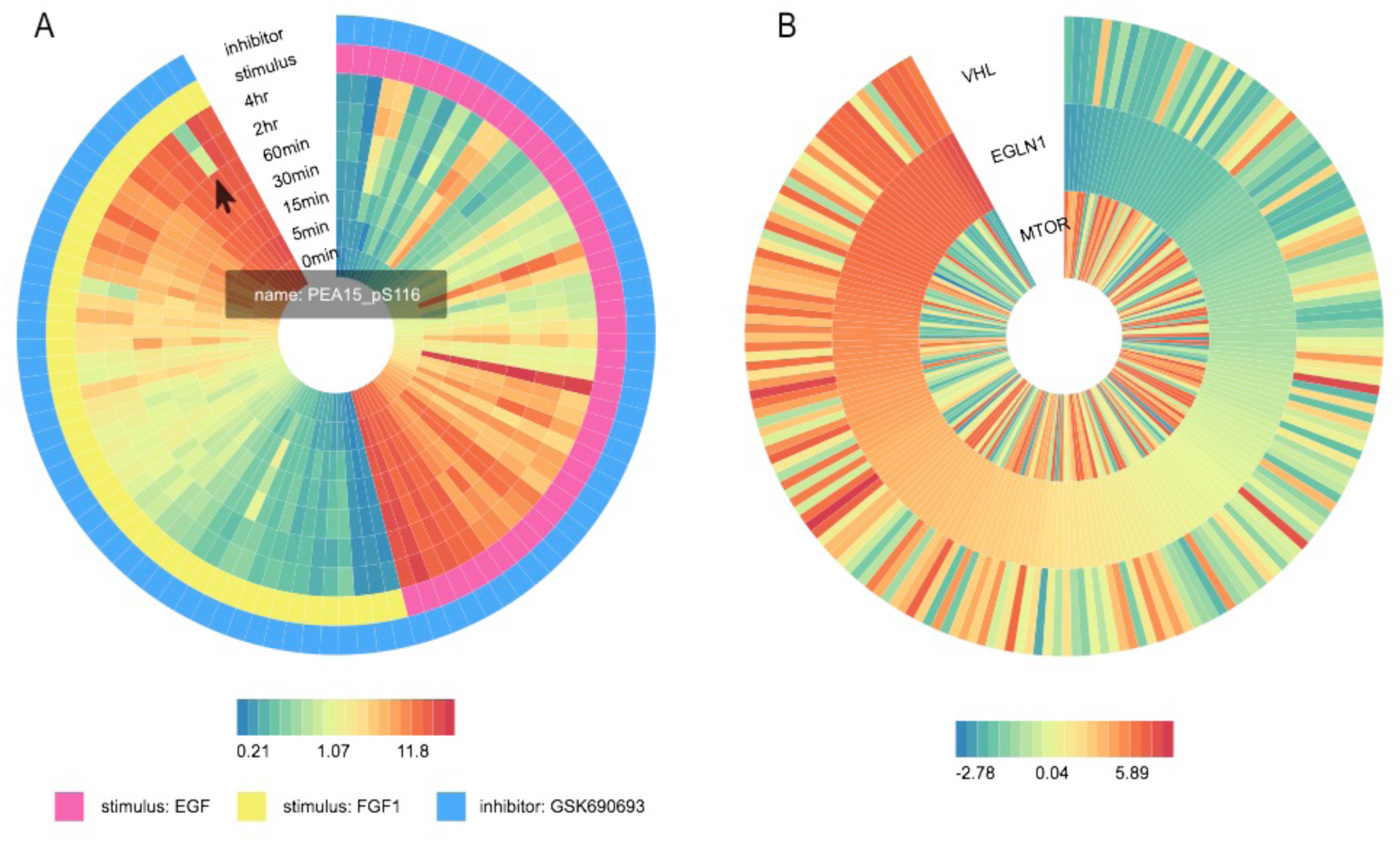
(A) Using the HPN-DREAM challenge data, the nested sorting of proteins by their expression levels at 0 min within each stimulus type reveals contrasting protein dynamics in response to different stimuli. (B) Using the AML challenge data, sorting of patients based on the EGLN1 expression indicates a positive correlation between VHL and EGLN1, and a partial negative correlation between EGLN1 and mTOR.

Interactive sorting can also provide insights of possible correlations between variables, especially those that only manifest in a sub-population of samples. To illustrate this utility, we applied Biowheel to visualizing data from the Acute Myeloid Leukemia (AML) Outcome Prediction Challenge^13^. This dataset consists of RPPA measurements of 231 proteins in 191 patients diagnosed with AML. As an example, Figure 2B shows the expression levels of EGLN1, VHL and mTOR from all patients, in which patients are sorted by the expression levels of EGLN1. It is clear to see a positive correlation in expression between EGLN1 and VHL across all samples in general, the finding of which is consistent with our existing knowledge about the roles of EGLN1 and VHL in the regulation of HIF1-α^14^. On the other hand, comparing the expression level of mTOR against that of EGLN1, we are able to see some negative correlation when the expression of EGLN1 is either low or high, but the trend is not consistent across all expression levels. A correlation analysis between EGLN1 and mTOR reveals a Pearson correlation coefficient of −0.310 (p-value = 1.3e-5) in all patients, −0.621 (p-value = 1.5e-4) in patients with an EGLN1 expression that is either above 1.0 or below −1.0, and −0.163 (p-value = 0.04) in patients with an EGLN1 expression that is between −1.0 and 1.0. Though mTOR was shown to regulate EGLN1 in previous experiments^15^, genetic aberrations and variations can complicate this relationship in AML^16^, as seen in this data. The association between EGLN1 and mTOR would be easily missed, if we directly conduct a correlation analysis across all samples without first visualizing the data. It is particularly important to look for mixed trends when analyzing cancer expression data, where heterogeneity is prevalent in both an intra-^17^ and inter-tumor^18^ manner.

Biowheel is a tool born from a community’s need to better visualize high-dimensional cancer expression data, and it is a design that was favored by our researcher peers in systems biology. As an effort to make powerful visualizations accessible to every biomedical professional including those without prior programming experience, we paid special attention to designing a user interface that is easy to learn and intuitive to use. We believe that Biowheel can be applied broadly to various research fields to improve and accelerate scientific discovery and sharing.

## METHODS

### Software

Biowheel was built based on the data-driven documents JavaScript library (d3.js)^11^, and is hosted at Amazon Web Services. The interface of Biowheel consists of three panels: an upper navigation bar for tasks at the system level (e.g. collecting feedback, login and help), a left panel for graphing option selection, and a main panel for visualization display. The accepted input file formats are csv and xlsx, and the recommend maximum file size is 1MB. Biowheel automatically recognizes numeric and non-numeric variables, thus missing data within continuous variables should be represented as empty cells in order for the software to correctly recognize. Once the files are uploaded, users will be prompted to select a sheet from the file for visualization. The uploaded data can be previewed and edited at any time. Selecting data for display consists of three components: selecting variables to display as rings (default is none), selecting samples to display as spokes by data filtering (default is selecting all samples), and selecting variables to display interactively as texts through tooltips. To customize the visualization, Biowheel offers options to modify colors, edit texts (e.g. displayed ring name), and position graphs. The final visualization can be exported into any of the three formats: svg, png and pdf.

### Data

The HPN-DREAM challenge data was first reformatted in order to present time points and experimental conditions as columns, and proteins as rows. The reformatted data from the BT20 cell line is provided in Supplementary Dataset 1, which includes a data subset that is sufficient to reproduce all the graphs presented in this paper (1^st^ sheet: simple) and the full data set (2^nd^ sheet: full). The AML Outcome Prediction Challenge data used in this study is available at https://www.synapse.org/#!Synapse:syn2501858.

## AUTHOR CONTRIBUTIONS

C.W.H., A.J.B. and A.A.Q. conceived of the Biowheel design. C.W.H. and A.J.B. implemented the software. C.W.H. wrote the manuscript. All authors reviewed and approved the final manuscript as submitted.

